# Functional properties of a disease mutation for migraine in Kv2.1/6.4 channels

**DOI:** 10.1101/2024.07.08.602400

**Authors:** Debanjan Tewari, Christian Sattler, Klaus Benndorf

**Author notes:** Correspondence should be addressed to D.T. or K.B.

## Abstract

Voltage-gated potassium (Kv) channels are integral to cellular excitability, impacting the resting membrane potential, repolarization, and shaping action potentials in neurons and cardiac myocytes. Structurally, Kv channels are homo or heterotetramers comprising four α-subunits, each with six transmembrane segments (S1-S6). Silent Kv (KvS), includes Kv5.1, Kv6.1-6.4, Kv8.1-8.2, and Kv9.1-9.3, they do not form functional channels on their own but modulate the properties of heteromeric channels. Recent studies have identified the Kv6.4 subunit as a significant modulator within heteromeric channels, such as Kv2.16.4. The Kv2.16.4 heteromer exhibits altered biophysical properties, including a shift in voltage-dependent inactivation and a complex activation. Current genetic studies in migraine patients have revealed a single missense mutation in the Kv6.4 gene. The single missense mutation, L360P is in the highly conserved S4-S5 linker region. This study aims to demonstrate the biophysical effects of the L360P mutation in Kv2.1 6.4 channels with a fixed 2:2 stoichiometry, using monomeric (Kv2.1/6.4) and tandem dimer (Kv2.1_6.4) configurations. Our findings suggest that the L360P mutation significantly impacts the function and regulation of Kv2.1/6.4 channels, providing insights into the molecular mechanisms underlying channel dysfunction in migraine pathology.

**Statement of significance:** This study elucidates the biophysical properties of the Kv6.4 L360P mutation, providing insights into its role in Kv2.1 6.4 channel function. Given the high conservation of the leucine residue in the Kv channel family and its association with migraine, our findings have significant implications for understanding the molecular basis of migraine pathophysiology. By analyzing channels with a fixed 2:2 stoichiometry, we highlight the impact of the L360P mutation on channel gating and inactivation. This research advances the knowledge of the silent Kv6.4 channel mechanism and its role in pathophysiology.

## Introduction

In humans, potassium ion channels are encoded by approximately 78 genes. Among these, the superfamily of voltage-gated potassium channels (Kv channels) is the largest, comprising about 40 genes in humans (1–3). Voltage-gated K^+^ channels play an important role in the excitability of different cell types, including various types of neurons and cardiac myocytes (2). They regulate the resting membrane potential and repolarization and shape the action potential and neuronal firing frequency (4–6). In the state of pain, some DRG neurons seem to employ potassium channels as a crucial mechanism to attenuate spontaneous activity (7).

Kv channels perform their biological roles by switching from closed to open states followed by inactivated states when the membrane potential is sufficiently depolarized (8, 9). Structurally, Kv channels are either homotetramers, formed by identical subunits, or heterotetramers, formed by homolog subunits. In homotetramers, four so-called α-subunits build the channels. Each subunit contains six transmembrane segments. The first four segments (S1–S4) comprise the voltage-sensing domains (VSDs), sensing the membrane potential. The positively charged residues in the S4 segment are the main voltage-sensing component in the VSD. The S5 and S6 transmembrane segments of each α-subunit arrange around the central axis to form the ion-conducting pore (10– 12). Upon membrane depolarization, the S4 segments move upwards via a combined rotating, tilting, and vertical displacement, which can be recorded as gating current in various types of Kv channels but also other voltage-gated channels (13, 14). These conformational changes are transmitted via an electromechanical coupling to an intracellular channel gate allowing channels to open. This intracellular gate is formed by the C-terminal ends of the four S6 transmembrane segments that obstruct in the closed channel the central ion-conducting pore via a bundle crossing (15).

Within the Kv superfamily, Kv1–4 subunits are known to combine to functional homotetramers. However, ten so-called silent subunits (KvS) cannot assemble into functional channels on their own. For mammals, these silent subunits are Kv5.1, Kv6.1–6.4, Kv8.1–8.2, and Kv9.1–9.3 (16). Though they cannot form own functional channels, they can be part of heteromeric channels and introduce special biophysical properties concerning homotetrameric channels formed by the included Kv1-4 subunits. This has been demonstrated e.g. by co-expressing Kv6.4 subunits with either Kv2.1 or Kv2.2 subunits (17). For example, Kv 2.1/6.4 heteromers show a shift of 40 mV in voltage-dependent inactivation to more negative potentials and a nearly five-fold reduction in current density compared to homomeric Kv 2.1 channels. Furthermore, voltage-dependent activation appears less steep than homomeric Kv 2.1 channels and the activation time course is more complex (18).

In many Kv channels, sustained depolarization induces a slow inactivation process that involves changes within the selectivity filter resulting in a non-conductive state (19). In some cases, slow inactivation can develop even before the opening of the intracellular channel gate, a process known as closed-state inactivation (9). The silent Kv 6.4 subunit has been reported to induce closed-state inactivation (CSI) in the heteromeric Kv2.1/6.4 channel (20). The molecular mechanism underlying the induction of CSI is still unknown. For Kv4 channels, pronounced CSI is well-characterized (21–23). CSI enables the channels to bypass the open state and become inactive. This allows the Kv4 channel to regulate the availability at the cell’s resting potential and, consequently, to modulate cellular excitability. Moreover, during repolarization, an inactivated channel also bypasses the open state and directly returns to the resting state, avoiding a potassium current (24, 25).

A recent study has highlighted a single mutation in the gene associated with Kv6.4 among migraine patients (26). A significant portion of these mutations is predicted to have detrimental effects. An example of such a mutation is L360P, located in the S4-S5 linker region of Kv 6.4. Notably, in the Kv-channel family, this leucine is highly conserved. Herein, we investigate the biophysical properties of the Kv6.4 L360P mutant in Kv2.1/6.4 channels with a fixed 2:2 stoichiometry by studying it either in a channel assembled from monomers (Kv2.1/6.4) or from linked tandem dimers (Kv2.1_6.4).

## Material Methods

### Molecular Biology

The genes for human Kv2.1 (KCNB1) and Kv6.4 (KCNG4) were inserted upstream of a T7 promoter within the pGEMHE vector, as described (27). By recombinant DNA cloning methods, a concatenated dimer of Kv2.1-Kv6.4 was engineered. The Kv2.1 subunit was amplified with primers containing a Kpn1 restriction site before the coding region and an Asc1 site behind an introduced linker sequence. The Kv6.4 subunit was amplified with primers containing an Asc1 site within a part of the linker sequence and an Xba1 site after the stop codon. The start codon in the second subunit and the stop codon in the first subunit were removed. The introduced linker sequence with ten amino acids is KARPTEGSLA. The mutations were generated with overlapping PCR protocols and subcloning of the respective fragments. The obtained clones were verified by restriction analysis and sequencing (Microsynth SEQLAB, Göttingen, Germany). cRNA for injection was prepared using the message mMACHINE T7 kit (Ambion Inc, Austin, USA). The quality of the cRNA was checked by gel electrophoresis.

### Electrophysiology and Data Analysis

Oocytes were procured from Ecocyte® (Castrop-Rauxel, Germany) or harvested from South African Xenopus laevis female frogs under an anesthetic condition with 0.1% tricaine (pH = 7.1; MS-222, Parmaq, Hampshire, UK). The oocytes were incubated in Ca^2+^ free solution (in mM: 82.5 NaCl, 2 KCl, 1 MgCl_2_, and 5 Hepes, pH 7.4) containing 3 mg/mL collagenase A (Roche, Grenzach-Wyhlen, Germany) for 105 mins. Then the oocytes of stages IV and V were selected and defolliculated. For the experiment, each oocyte was injected with 30 nL of 0.2 microgram/microliter of mRNA. Further, injected oocytes were kept at 18°C for 1 day in Barth’s medium for the synthesis and trafficking of ion channels to the plasma membrane. The whole-cell current of Kv 2.1, 6.4, and other concatemeric constructs was measured from oocytes using a two-electrode voltage clamp technique (725C amplifier, Warner Instruments, LLC, Hampden, MA, USA) at room temperature. Glass electrodes (Brand GmbH + Co KG, Wertheim, Germany) were filled with 3 M KCl solution. The resistance of the glass electrodes was between 0.3 – 0.9 MΩ. The bath solution contained 96 mM NaCl, 2 mM KCl, 1.8 mM CaCl_2_, 1 mM MgCl_2_, and 10 mM HEPES, pH = 7.4. For activation currents were triggered by test potentials from −100 to +100 mV in 10 mV increments from a holding potential of −90 mV. The duration of test potentials was 300 ms as mentioned in the figures. To study inactivation currents were triggered by test potentials from -120 mV to +60 mV in 10 mV increments from a holding potential of – 90 mV. The duration of the test potential was 4s and the intermediatory pulse was 10-15 s. In the case of both activation as well as inactivation data were collected at a sampling frequency of 5.00 kHz. For data acquisition and preliminary analysis, we employed Patchmaster software (HEKA Elektronik GmbH, Reutlingen, Germany). The steady-state activation was determined from the amplitude of the instantaneous tail currents at +30 mV. The experimental data points measured at different voltages fitted with a single Boltzmann’s equation: I/I_max_=A/{1+exp[zδF(V-V_h1_)/RT]} where V_h1_ is the midpoint voltage of half-maximum activation, V is the membrane potential, zδ is the charge and I_max_ is the maximum current.

## Results

### Functional characterization of both homomeric Kv 2.1 and heteromeric Kv2.1/6.4 channels

First, we investigated the influence of Kv6.4, a member of the KvS family, on the biophysical characteristics of Kv2.1 regarding steady-state activation and inactivation. We first conducted experiments where the mRNA of Kv2.1 and Kv6.4 constructs was mixed in a 1:4 ratio and co-expressed the subunits in *Xenopus* oocytes. Currents were recorded using a pulse protocol outlined in Figures 1A and E. While analyzing the activation curve, a noticeable shift in both V_1/2a_ and the slope factor was observed (for quantifying the changes, the activation curve was fitted with a Boltzmann equation). For Kv2.1 alone, the V_1/2a_ value was -1.1 ± 0.5 mV, quite consistent with previous reports (18, 28). The slope factor was 13.6 ± 0.3 mV as shown in Figures 1C and D. Nonetheless, when Kv 6.4 was co-expressed with Kv2.1, there was a clear change in V_1/2a_. V_1/2a_ for the co-expressed construct was -15.5 ± 0.6 mV and the corresponding slope factor was 17.4 ± 0.6 mV (Figure 1). We also investigated inactivation for both Kv 2.1 and Kv 2.1/6.4 constructs when co-expressed with the RNA in the mentioned 1:4 ratio. The inactivation protocol is depicted in Figure 1E. A significant shift in the inactivation curve was observed for Kv2.1/6.4 compared to the Kv 2.1 homomer. For Kv 2.1, V_1/2i_ of the inactivation curve was -21.0 ± 0.5 mV, whereas Kv2.1/6.4 exhibited a V_1/2i_ of -58.3 ± 1.1 mV (Figure 1F). These values were determined by fitting the data points with a single Boltzmann equation. This shift in inactivation confirms the assembly of the Kv 6.4 subunits with Kv 2.1. Hence, both subunits express as heteromers when injected mRNA in the 1:4 ratio.

**Figure 1.**
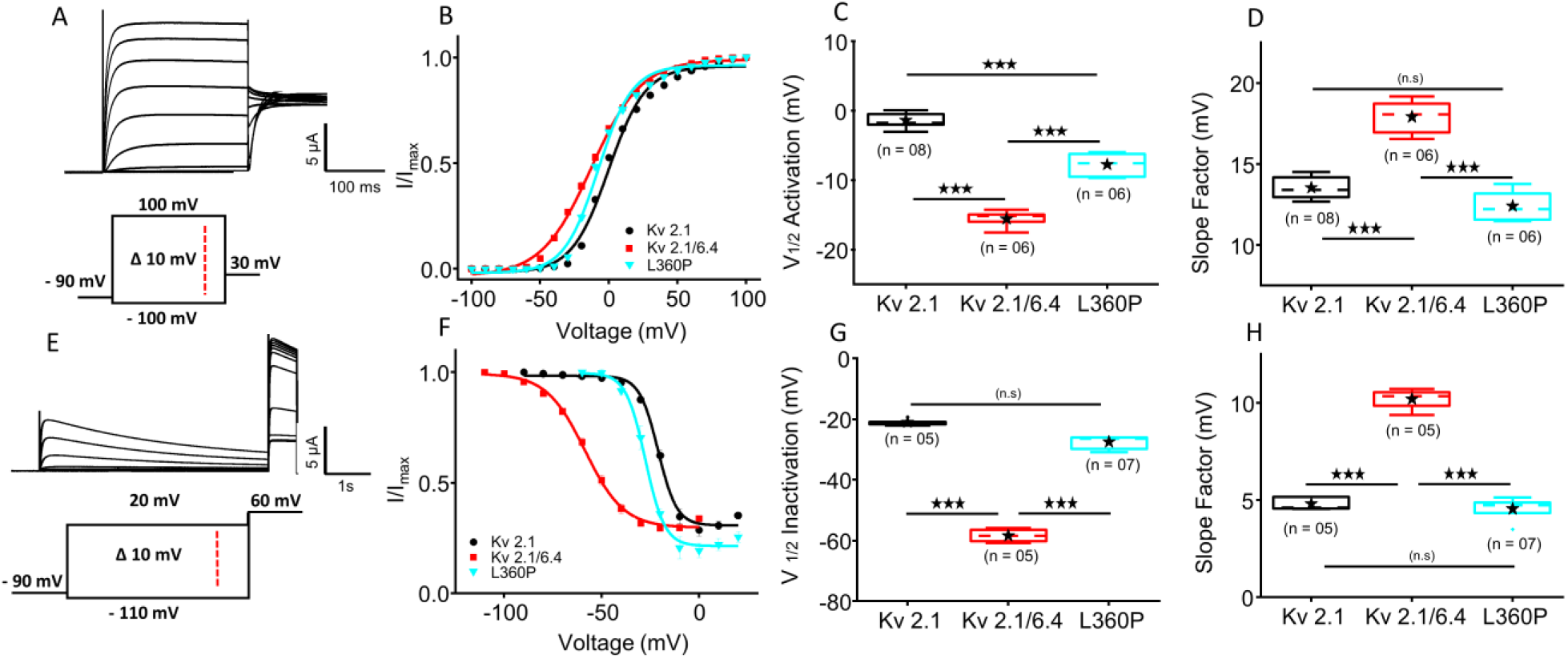
Voltage-activation relationships for homo and heteromeric Kv channels. **(A)** Representative current traces of Kv 2.1 activation. **(B)** Steady-state activation curves were derived from tail currents at +30 mV for Kv 2.1 (black), Kv 2.1/6.4 (red), and Kv 2.1/6.4L360P (cyan, labeled as L360P in the figure). Data are presented as mean ± SEM. **(C and F)** Box plots showing half-activation voltage (V_1/2a_) and slope factor, obtained by fitting a single Boltzmann equation. A black star inside each box indicates mean values. **(E)** Representative current traces demonstrating the inactivation of Kv 2.1 with the pulse protocol depicted below. **(F)** Steady-state inactivation curves obtained from tail currents at +60 mV for Kv 2.1 (black), Kv 2.1/6.4 (red), and Kv 2.1/6.4L360P (cyan, labeled as L360P in the figure). Data are presented as mean ± SEM. **(G and H)** Box plots showing half-inactivation voltage (V_1/2i_) and slope factor, obtained by a fit with a single Boltzmann equation. Mean values are indicated with a black star inside each box. For all values statistical significance (*** p < 0.001, n.s. for not significant) was determined using one-way ANOVA followed by post hoc Tukey test.

Subsequently, a specific mutation was introduced into the Kv 6.4 subunit. The mutation lies within the S4-S5 linker of the Kv 6.4 subunit and the structure of Kv 6.4 was visualized and colored using UCSF Chimera (v.1.7.1) (29). It is noteworthy to mention that the leucine residue at this position is highly conserved not only in Kv6 subunits but also across Kv channels in general (Figure 2). In this context, the specific mutation L360P in Kv6.4 is of particular interest due to its association with headaches in migraine patients. To understand the impact of this mutation on channel behavior, the Kv6.4L360P mutant was co-expressed with Kv 2.1 and its activation and inactivation properties were measured (Figures 1B and F). Pronounced changes were observed in both activation and inactivation compared to the Kv 2.1/6.4. The activation curve exhibited a steeper slope compared to Kv2.1/6.4 (Figure 1D and H). This suggests that the mutant L360P in Kv6.4 affects the voltage dependence of channel activation, making it more sensitive to changes in membrane potential. In contrast to the markedly left-shifted inactivation curve of Kv2.1/6.4, the inactivation curve of the Kv 2.1/6.4L360P channel shifted towards the right. These findings suggest that the L360P mutation in the Kv6.4 subunit significantly impacts the functional properties of Kv2.1/6.4 channels, altering both activation and inactivation.

**Figure 2.**
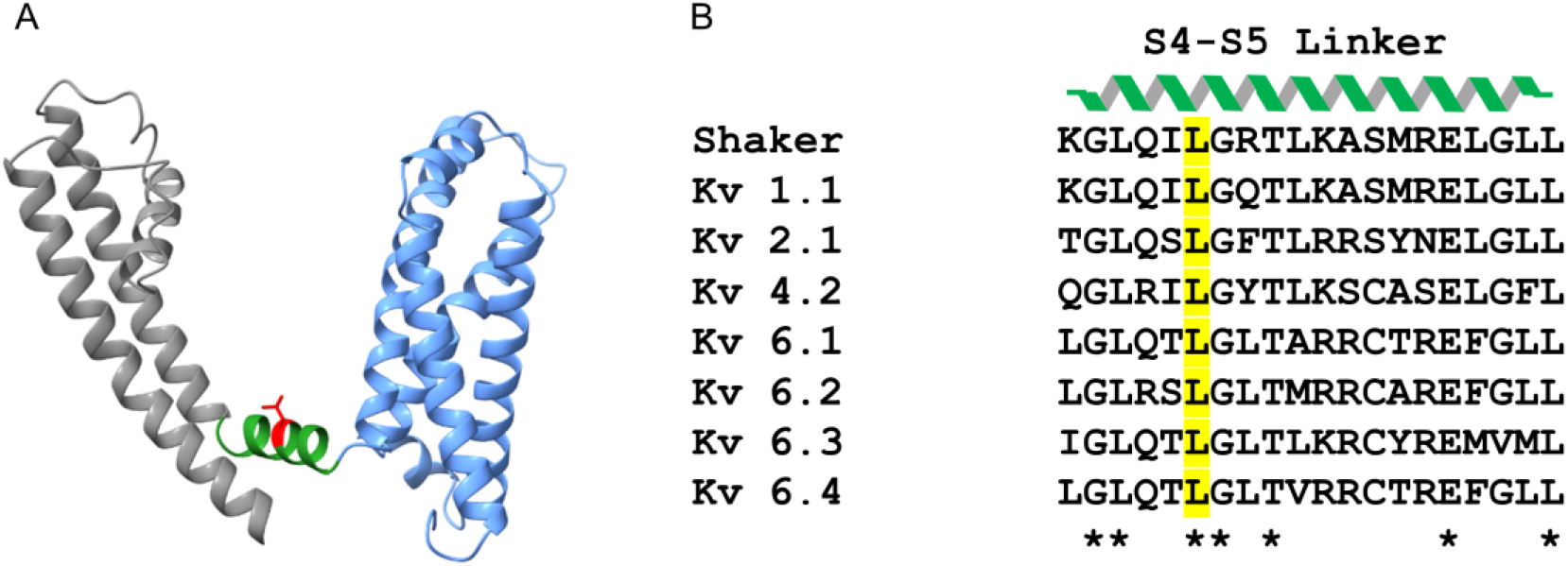
Cartoon of a Kv 6.4 subunit. **(A)** Structure of a Kv 6.4 subunit downloaded from Alpha Fold Protein Structure Data Base (UniProt-Q8TND1). The voltage-sensing domain (VSD) is highlighted in corn blue, with the S5 and S6 domains in dark grey. The S4-S5 linker is depicted in green, and the leucine residue is shown in red. (B) The sequence alignment of the S4-S5 linker highlights the highly conserved leucine residue at position 360 in yellow.

### Biophysical properties of concatenated Kv 2.1_6.4L360P

Recent research has revealed a fascinating aspect of Kv 2.1 potassium channels when expressed with Kv 6.4. They express in a consistent 2:2 ratio with Kv 6.4 subunits. Markedly, the arrangement of Kv6.4 subunits within this complex follows a specific pattern - they never appear in adjacent positions (18) but consistently occupy diagonally opposite positions (Figure 3A). To maintain both the stoichiometry and the consistent subunit orientation, we have linked Kv2.1 and Kv 6.4 subunits together using an engineered 10-amino acid linker. This molecular manipulation ensures that the 2:2 ratio and the diagonal arrangement of Kv6.4 subunits are maintained. After fusing the Kv2.1 subunit with the mutated Kv6.4L360P variant, the resulting heteromeric constructs Kv2.1_6.4L390P were thoroughly analyzed with the protocol outlined in Figure 1. Notably, there was no detectable difference in activation and inactivation between co-expressed Kv2.1/6.4 and concatenated Kv2.1_6.4 channels (Figures 1B, F and Figure 3B, C).

**Figure 3.**
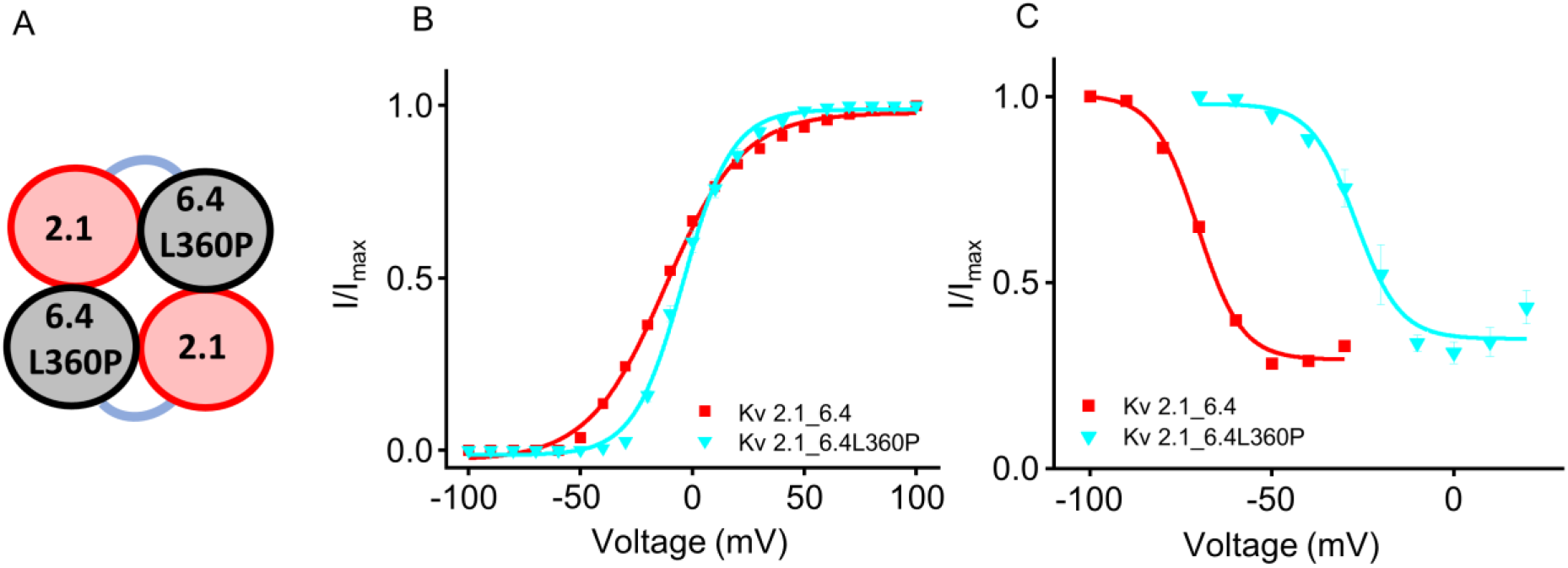
Biophysical characteristics in the dimeric form for Kv 2.1_6.4 and 2.1_6.4L360P. **(A)** Cartoon diagram of the concatenated Kv 2.1_6.4L360P in 2:2 ratio with diagonally opposite subunit arrangement. **(B)** Steady-state activation was obtained from the tail current at +30 mV for 2.1_6.4 (red) and Kv 2.1_6.4L360P (cyan) using the protocol as shown in Figure 1. **(C)** Steady-state inactivation measured from the tail current at +60 mV for Kv 2.1_6.4 (red) and Kv 2.1_6.4 L360P (cyan) using the protocol depicted in Figure 1. For activation as well as inactivation data are presented as mean ± SEM.

When the mutant associated with the headache condition was evaluated within the dimeric construct Kv2.1_6.4L390P, notable differences emerged in V_1/2_ and the slope factor (Table 1). Activation showed a steeper slope and inactivation was shifted towards depolarizing potentials when compared to the Kv2.1_6.4 construct. The monomeric channels Kv2.1/6.4L360P and the dimeric channels Kv2.1_6.4L360P showed a closely similar result, suggesting that the mutated monomer is incorporated in a similar way.

**Table 1.** Biophysical properties of concatenated Kv 2.1_6.4 and Kv 2.1_6.4L360P.

| **Constructs** | **V<sub>1/2</sub>** | **Slope factor** |
| --- | --- | --- |
| **Activation** |  |  |
| Kv 2.1_6.4 (n= 5) | - 13.9 ± 0.8 | 16.7 ± 0.2 |
| Kv 2.1_6.4L360P (n= 6) | - 4.0 ± 1.1 | 11.4 ± 0.4 |
| **Inactivation** |  |  |
| Kv 2.1_6.4 (n= 7) | - 66.2 ± 0.8 | 12.2 ± 0.6 |
| Kv 2.1_6.4L360P (n= 6) | - 24.8 ± 1.3 | 5.8 ± 0.2 |

Values are given as mean ± SEM and n is the number of oocytes analyzed. The mid points of activation and inactivation (V<sub>1/2</sub>), represented in millivolts, and the slope factor were obtained from a single Boltzmann fit. For all values statistical significance was determined using student t test (p value less than 0.001).

### The L360P in Kv6.4 rescues the heteromeric channel from closed-state inactivation

The Kv6.4 subunit is known to shift voltage-dependent inactivation in heteromeric Kv2.1/6.4 channels to hyperpolarized potentials by approximately 40 mV. Additionally, it induces closed-state inactivation within the heteromeric channel. Notably, the mutant Kv6.4L360P induces a shift in inactivation towards depolarizing potentials, approximating its behavior more closely to that in WT Kv2.1. We therefore considered the recovery process from closed-state inactivation in the Kv6.4L360P mutant by determining the recovery rate (see protocol in Figure 4A). Initially, a control pulse of +60 mV was administered to gauge the initial current amplitude for 100 ms. Then a 400-millisecond pulse to -130 mV was applied, to facilitate the recovery of all channels from inactivation, followed by a 5-second pulse to -60 mV to induce a certain level of closed-state inactivation. Finally, a pulse of variable duration to -90 mV allowed the channels to recover from inactivation, followed by a second test pulse to +60 mV. I/I_max_, indicating the recovered proportion of channels from their inactivated state, was plotted against the time spent at the recovery pulse to -90 mV (Figure 4C). The results revealed that the Kv2.1/6.4 channel exhibited a recovery rate of 192.9 ± 19.2 milliseconds (n=6). In contrast, the Kv2.1/6.4L360P mutant displayed a markedly longer recovery rate of 1474.6 ± 242.0 milliseconds (n=5; Figure 4D). This effect underscores a notable difference in the behavior of the two constructs. Hence, the L360P mutation in the S4-S5 linker of Kv6.4 influences the channel’s response strongly, rescuing it from closed-state inactivation.

**Figure 4.**
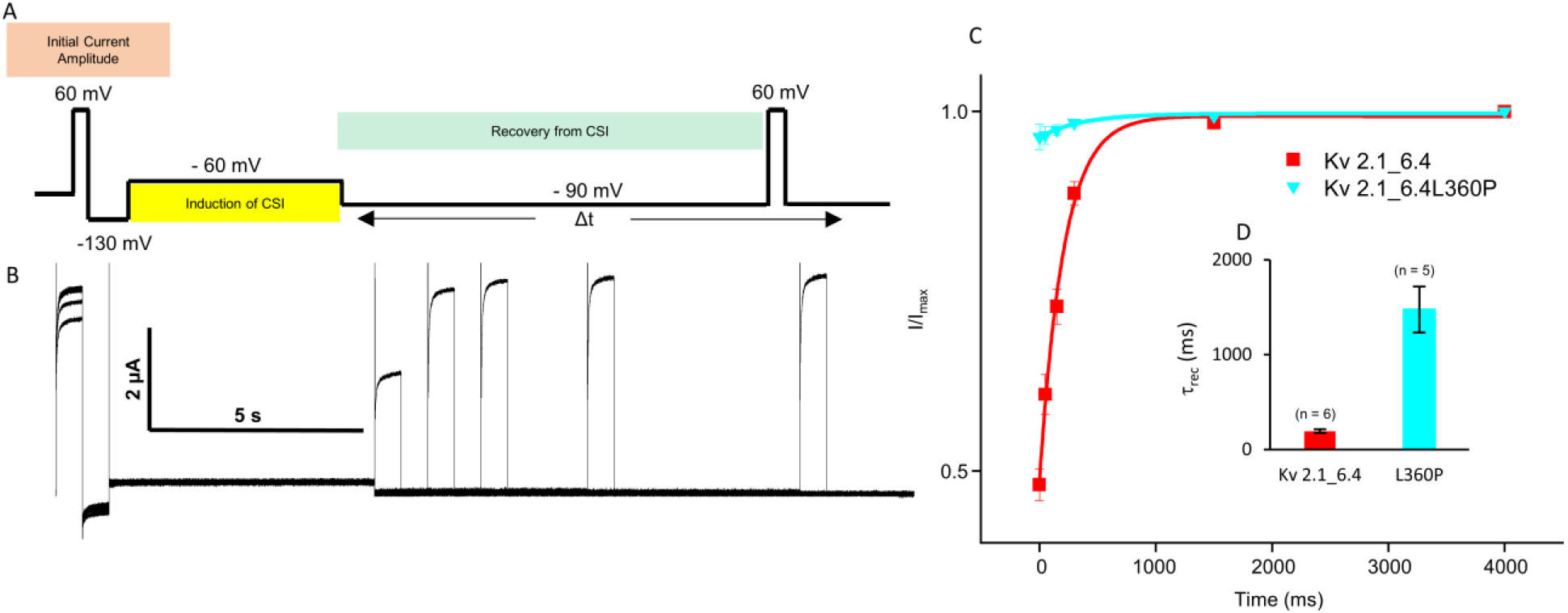
L360P rescues the channel from closed-state inactivation. (**A and B**) Representative current recordings were obtained using the voltage pulse protocol shown at the top to study the recovery of Kv 2.1_6.4 from closed-state inactivation (details are provided in the results section). **(C)** Recovery of Kv2.1_6.4 dimer and Kv2.1_6.4L360P from closed-state inactivation was analyzed by plotting the normalized I/I_max_ current amplitudes at +60 mV as a function of the pulse duration at the −90 mV voltage pulse. Single exponential functions were fitted to the data for the Kv2.1_6.4 dimer and Kv2.1_6.4L360P (red and cyan, respectively). **(D)** Recovery time constants for the closed-state inactivation of Kv2.1_6.4 (red) and Kv2.1_6.4L360P (cyan, labeled as L360P in the figure). The L360P mutation rescued the recovery of Kv2.1_6.4 from closed-state inactivation (n = number of cells).

## Discussion

This study aimed to investigate the biophysical properties of Kv2.1 potassium channels when coexpressed with the Kv6.4L360P subunit. The implications of our findings seem to be relevant to understanding the pathophysiology of migraine headache associated with this mutation in Kv6.4. Regarding the role of the Kv6.4 subunit in Kv2.1/Kv6.4 channels, our results confirm and extend previous observations about the modulatory role of Kv6.4. Consistent with prior research, Kv6.4 does not form functional channels by itself but significantly alters the characteristics of Kv2.1 channels (20, 28, 30, 31). Introducing the mutation L360P into the S4-S5 linker of Kv6.4, as identified in the context of the pathogenesis of migraine, brought about additional changes in channel behavior when co-expressed. The mutation led to a rightward shift of inactivation while activation remained unaffected, suggesting a decreased propensity for the channel to enter closed-state inactivation (32). One of our key findings is the dramatically altered recovery kinetics from closed-state inactivation: it is strongly slowed with the L360P mutant compared to the wild-type co-expression. The L360P mutant in the Kv 6.4 subunit, when co-expressed with Kv2.1, exhibits electrophysiological properties similar to the wild-type Kv2.1. Besides activation, steady-state inactivation of these channels is shifted to the right, approximating that of WT Kv2.1 channels. This shift means that more channels are recused from closed-state inactivation, resulting in more active channels.

Generally, the function of Kv2 channels in neurons is very substantial, influencing action potential width and strongly regulating repetitive firing (33–35). Repetitive depolarization accelerates Kv channel inactivation, followed by a broadening of action potential, thus limiting the firing rate. The incorporation of Kv6.4 subunits into Kv2.1 channels results in a higher proportion of inactivated states because of CSI. This fine-tuning may limit the action potential rate and the excitability. The loss of this modulatory function in Kv6.4 L360P mutant channels might contribute to the pathophysiological basis of migraine with increased excitability in respective neurons.

Previous reports indicate that in most of the Kv channels, there is a dynamic molecular interaction between the carboxy-terminal end of S6 (referred to as S6c) and the S4-S5 linker (23, 25, 36). This interaction plays an important role in the electromechanical coupling, translating the movement of the VSD into the opening of the bundle crossing (BC) gate (37, 38). In case, the S4-S5 linker and S6c interface become uncoupled, it disrupts the normal process of BC gate opening. As a result, the channel is unable to transit into a conductive state, leading to a non-conductive or inactivated state. The opening of the BC gate in shaker-related Kv channels relies on the subunit co-operativity (39). Even if one subunit, like Kv 6.4 in a heteromeric channel, has its electromechanical coupling disrupted, it can cause the channel to become non-conductive or enter an inactivated state (20). A similar uncoupling mechanism has been demonstrated in hyperpolarized cyclic nucleotide-gated (HCN) channels. In these channels, the uncoupling between the voltage-sensing domain (VSD) and the channel gate takes place at hyperpolarizing potential leading to desensitization to voltages observed as gating change immobilization (40). Intriguingly, similar desensitization to voltages occurs in Kv 4.2 channels, which is displayed as closed-state inactivation (41). Leucine 360 is positioned near the N terminal of the S4-S5 linker within potassium (Kv) channels. It is important to mention that this amino acid is highly conserved across diverse Kv channels, suggesting functional importance. In case of a mutation replacing Leucine by Proline, this might cause the S4-S5 linker to adopt a different conformation which can alter the molecular interactions between the S4-S5 linker and S6c. One possible consequence of this structural change might be the prevention of a non-conducting state reached by CSI before the pore opening.

## Author contributions

D.T. did the measurements, analyzed the data, and wrote the manuscript. C.S. did the molecular biology. K.B. supervised the study and edited the manuscript.

## Acknowledgment

S. Bernhardt, C. Ranke, U. Singer for excellent technical assistance. The work was supported by the Jena University Hospital.

## Note

**While our manuscript was under review there was a submission in bioRxiv with a similar L360P mutation in the Kv 6.4 related to migraine by Lacroix et al. The paper titled “KCNG4 GENETIC VARIANT LINKED TO MIGRAINE PREVENTS EXPRESSION OF KCNB1.”**

